# Cockayne Syndrome B protein selectively interacts and resolves intermolecular DNA G-quadruplex structures

**DOI:** 10.1101/2021.03.25.436565

**Authors:** Denise Liano, Marco Di Antonio

## Abstract

Guanine-rich DNA can fold into secondary structures known as G-quadruplexes (G4s). G4s can form from a single DNA-strand (intramolecular) or from multiple DNA-strands (intermolecular), but studies on their biological functions have been often limited to intramolecular G4s, owing to the low probability of intermolecular G4s to form within genomic DNA. Herein, we report that the endogenous protein Cockayne Syndrome B (CSB) binds with picomolar affinity to intermolecular G4s, whilst displaying negligible binding towards intramolecular structures. We also observed that CSB can selectively resolve intermolecular G4s in an ATP independent fashion. Our study demonstrates that intermolecular G4s formed within ribosomal DNA are natural substrates for CSB, strongly suggesting that these structures might be formed in the nucleolus of living cells. Given that CSB loss of function elicits premature ageing phenotypes, our findings indicate that the interaction between CSB and ribosomal DNA intermolecular G4s is essential to maintain cellular homeostasis.

## Introduction

Cockayne Syndrome (CS) is a rare autosomal-recessive disorder, characterised by premature ageing, UV-sensitivity, progressive neurological degeneration and growth failure^1^. Loss of function mutations in *CSB* (also known as *ERCC6*) gene have been associated with approximately 65% Cockayne Syndrome cases^2^. Experimental evidence revealed the importance of CSB in chromatin remodelling and in transcription-coupled DNA repair (TCR) mechanisms^3,4,5^. Beside its nucleoplasm localization, immunofluorescence and immunoprecipitation studies have demonstrated nucleolar localization of CSB^6^. Nucleoli are non-membrane enclosed, highly conserved, sub-organelles within the nucleus where transcription and ribosomal RNA (rRNA) processing occurs^7^. Recent studies have revealed that CSB is key to promote efficient rRNA synthesis^8,9^. Due to the guanine-rich nature of human rDNA, G-quadruplex DNA secondary structures (G4s)^10,11^ have been postulated to form at the rDNA level^12^, which has been recently reinforced by the direct visualisation of G4s structures formation in living human under non-perturbative conditions^13^. Previous studies have suggested that rDNA transcription might promote the formation of these secondary structures^14^ and transcriptional stalling at rDNA G4s observed in cells lacking CSB activity has suggested a role of CSB in resolving rDNA G4s^15^. Importantly, the increased variability in rDNA transcriptional rate with age around G4 regions is linked to ageing features, including hyperactivation of poly (ADP-ribose) polymerase 1 (PARP1), nicotinamide adenine dinucleotide (NAD^+^) depletion and mitochondrial dysfunctions^15,16,17^. Altogether, these findings have suggested that accumulation of unresolved G4 structures within rDNA could promote ageing phenotypes, which are exacerbated in Cockayne Syndrome patients that lack functional CSB. The link between G4s and CSB was further corroborated by the observation that treatment with the G4 ligands Pyridostatin (PDS^18^) and CX-5461^19^ in *C. elegans* models could recapitulate the premature ageing phenotype observed in CSB^-/-^ cells^15^. This hypothesis was recently reinforced by confocal imaging using a fluorescent-tagged CSB protein (CSB-GFP) that have revealed how treatment with CX-5461 could displace CSB from the nucleoli of human living cells^6^, suggesting that the ligand could outcompete CSB for binding to rDNA G4s. On the other hand, biochemical experiments on the resolution of rDNA G4 structures by CSB actually revealed that this protein is a modest G4-helicase^15^, which is surprising given the strong transcriptional stalling at rDNA G4s observed in CSB impaired cells. This highlighted that certain structural features somewhat exclusive to rDNA and to the nucleolus environment make CSB a selective interactor of rDNA G4s and that such binding might be key to the premature ageing phenotype, as observed in Cockayne Syndrome patients lacking functional CSB or in PDS/CX-5461 treated model organisms.

Herein, we investigated the interaction between both recombinant full length CSB (CSB-FL) and its DNA binding domain (CSB-HD) towards a panel of G4-forming sequences, with particular interest on G4 structures formed within rDNA. We observed that CSB is able to resolve only a small sub-population of G4s formed by rDNA. Further analysis revealed that CSB has an ATP-independent G4-resolvase activity that is limited to intermolecular G4s^20^ that can be formed by rDNA sequences, whilst being inactive towards intramolecular G4s formed by both rDNA and non-rDNA sequences. We further observed that CSB can bind with astonishing picomolar affinity intermolecular rDNA G4s, whilst displaying negligible binding towards unimolecular G4s formed by the same rDNA sequences. The unprecedented selectivity for intermolecular G4s displayed by CSB suggests that these structures are likely to be formed under physiological conditions at rDNA level, possibly promoted by the high-density environment typical of the nucleoli and the high GC richness of rDNA. We further demonstrated that treatment with the G4 ligands PDS or CX-5461 can compete out CSB bound to rDNA intermolecular G4s, supporting a model by which the nucleolar localisation of the protein is mediated by selective interaction with intermolecular G4s, and that abrogation of such interaction might be responsible of the premature ageing phenotypes observed in previous studies. To the best of our knowledge this is the first example of an endogenous protein capable of discriminating intermolecular from intramolecular G4-structures with high selectivity, suggesting differential biological functions and protein recognition of G4s based on their forming stoichiometry.

## Results

### CSB is not an efficient G4-resolvase with limited activity towards rDNA

The ability of CSB to resolve G4 structures formed by a specific rDNA sequence that has been recently reported^15^, prompted us to systematically investigate CSB as a G4-helicase against a small panel of G4-forming sequences. We therefore expressed and purified the full-length protein (CSB-FL: 1-1493, Fig. 1a, b) from *S. frugiperda* (*Sf9*) using a previously established protocol^21,22^. Because the DNA binding domain of CSB contains a DEGH box that is typical of common helicases, we decided to also express this helicase-like domain^23^ (CSB-HD: 498-1002 aa, Fig. 1a, c) from *E. coli* cells to explore if this domain alone was able to resolve G4 structures. We firstly assessed the ability of CSB-FL to resolve rDNA G4s in a gel-based helicase assay (see Fig. 1d). We initially focussed our attention on a specific rDNA G4 sequence (see Supplementary Table S1 for all the sequences used in this section), as this was the only G4 substrate for CSB previously reported^15^. To assess the G4-helicase activity, we incubated the recombinant CSB protein with the rDNA G4 forming sequence, annealed in KCl, in the presence of its complementary strand for increasing incubation times (from 0.5 to 40 minutes).

**Figure 1.**
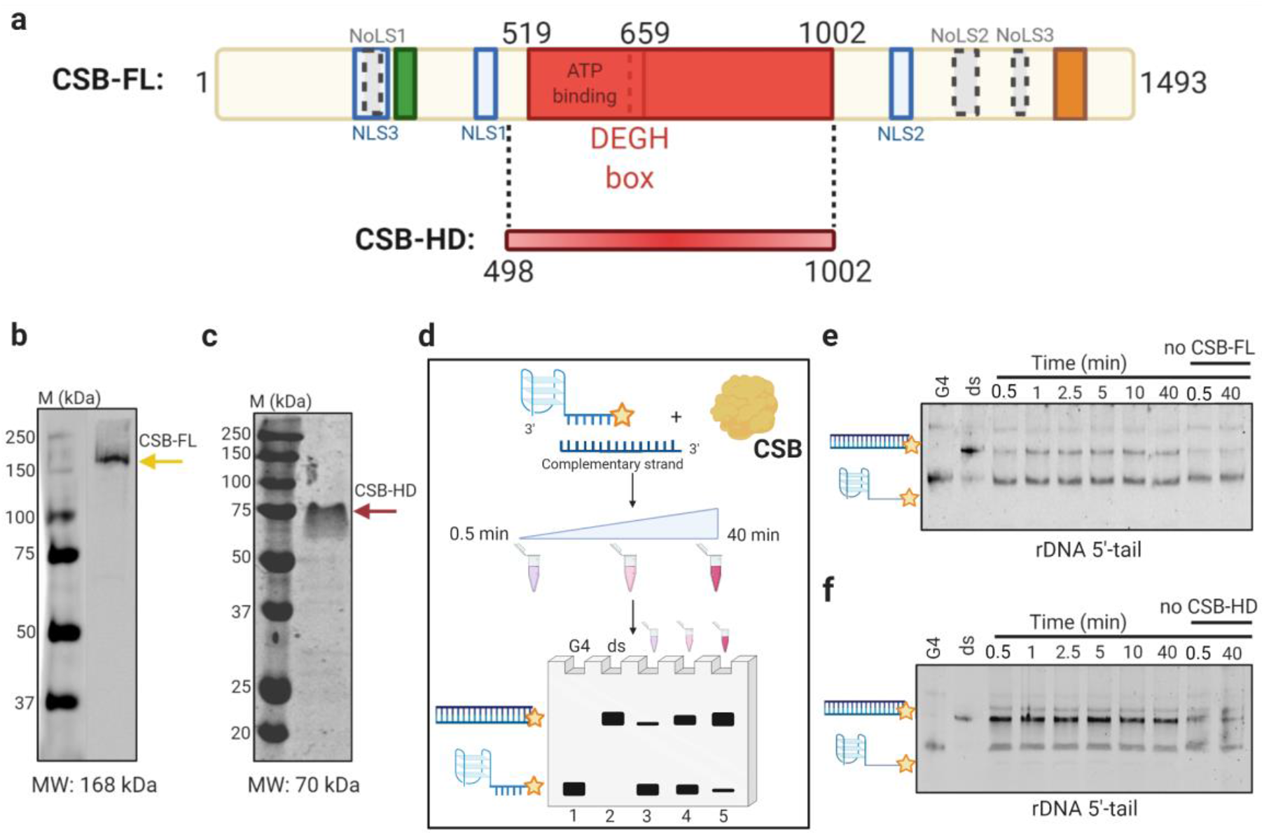
CSB has specific rDNA-G4 resolvase activity. (**a**) Schematic representation of the CSB-FL (1-1493 aa) and the isolated CSB-HD (498-1002 aa). Different regions of CSB-FL are indicated with different colours. The three nuclear and nucleolar localisation sequences are indicated as light blue and grey dashed boxes, respectively. The helicase-like domain (519-1002 aa) is represented in red and contains the ATP-binding domain and the DEGH box. The most stable CSB-HD construct that we tested (498-1002 aa) is reported at the bottom in a gradient-red line. (**b**) Western blot (WB) gel using anti-CSB antibody confirmed the presence of the CSB-FL (168 KDa, yellow arrow). (**c**) SDS-page gel indicating the presence of the CSB-HD. The red arrow indicates the expected molecular weight (MW) of CSB-HD (MW: 70 KDa). (**d**) Scheme of the gel-based helicase assay used in this work. A Cy5-fluorophore labelled 5’-tail G4-forming oligonucleotide was incubated with its complementary strand in presence or absence of CSB (in yellow) for increasing times (0.5 to 40 minutes). The products were then separated on a polyacrylamide gel where is possible to identify double stranded (ds) products as bands at the top of the gel (pre-annealed ds control: lane 1) and unresolved G4 bands at the bottom of the gel (G4 control: lane 2). The intensity of the G4 bands decrease upon G4-resolution and ds formation with consequent increasing of the intensity of the ds bands (lanes 3 to 5). (**e**) Helicase assay gel using 5’-tailed rDNA G4 after incubation with or without CSB-FL. (**f**) Helicase assay gel using 5’-tailed rDNA G4 after incubation with or without or CSB-HD.

Specifically, the rDNA G4 sequence used was labelled at its 5’-end with a Cyanine 5 (Cy5) fluorophore and included a 5’-20bp tail for consistency with previous reports^15^. The products were then separated on a polyacrylamide gel, confirming that CSB was able to promote the resolution of the rDNA G4 tested in the absence of ATP, as assessed by the formation of a new band that could be ascribed to the double stranded (ds) product (Fig. 1e). However, we noticed that CSB-FL displays only a modest G4-helicase unfolding activity even after 40 minutes incubation, with most of the substrate still present in the form of a folded G4 (Fig. 1e). Indeed, the gel quantification revealed a statistically significant increase (p=0.02 using a two-tailed Student’s t-test^24^, see Supplementary Fig. S1a and Supplementary Table S2 for all the plotted data) of the percentage of ds product formed after incubation (0.5 to 40 min) with CSB (24.6%) compared to the control without the protein (8%), but the G4 resolution remains partial and, most importantly, does not increase with increasing incubation times (Fig. 1e), suggesting that only a subpopulation of the G4 structures present is efficiently unfolded whilst most of the substrate remains unaltered.

To further confirm the ability of CSB to resolve G4 structures formed within rDNA, we tested a different 5’-tailed rDNA-2 sequence which was also reported to fold in a G4 structure^12^ (see Supplementary Table S1). We measured an increasing G4-resolution in presence of CSB (Supplementary Figs. S1b, S2a) and, similarly to what observed with the first sequence tested, only a partial 15% (Supplementary Table S2) increase in dsDNA formation was measured after exposure to the protein. To evaluate if the presence of a 5’-tail was essential for G4-resolution, we also tested untailed rDNA (see Supplementary Table S1). Interestingly, negligible difference on dsDNA formation was observed after incubation of this substrate with CSB (Supplementary Figs. S1c, S2b, Supplementary Table S2), suggesting that similarly to other G4- and dsDNA-helicases, CSB requires a single strand overhang tail in order to initiate the unwinding^25,26^. We next decided to test non-rDNA G4s to assess whether such modest helicase activity could also be observed for other G4-forming sequences or if it was limited to the rDNA substrates (see Supplementary Table S1 for the sequences used in this section).

Notably, no G4-resolvase activity promoted by CSB-FL was observed for all the panel of G4s tested, including c-KIT1, hTELO, HRAS and c-MYC functionalised with the same 5’-tail sequence used for the rDNA G4s (Supplementary Figs. S1d, e, f, g, S2c, d, e, f, Supplementary Table S2). Similarly, we failed to observe G4-resolvase activity on both 3’-tailed or untailed c-MYC (Supplementary Figs. S1h, i, S2g, h, Supplementary Table S2) and on untailed c-KIT1 (Supplementary Figs. S1l, S2i, Supplementary Table S2) substrates. To confirm the correct detection of dsDNA products from G4 substrates, we tested a mutated 5’-tailed c-MYC (c-MYC-mut 5’-tail) sequence as control that is no longer able to form G4. As expected, we observed complete ds formation under identical conditions (∼70%) with negligible difference observed when CSB was added (Supplementary Figs. S1m, S2l, Supplementary Table S2), suggesting that simple DNA pairing between the two complementary strands is observed under these conditions.

### CSB DNA binding domain is responsible for the observed helicase activity

We then asked whether the DNA binding domain of CSB was sufficient to recapitulate the helicase activity observed with the full-length CSB protein on rDNA G4 substrates. To achieve this, we performed a multiple sequence alignment (Clustal Omega, EMBL-EBI^27,28^) to select the most conserved region referred to the helicase-like domain of the protein between a wide range of different organisms (Supplementary Fig. S3a). Then, we performed a secondary structure prediction to ensure that in the folded protein there was no destruction of helices (using PSIPRED 4.0^29^, Supplementary Fig. S3b), which could affect the stability of the protein. The conserved sequence (CSB-HD: 498-1002, Fig. 1a) was therefore cloned into a pCS46_6xHistidine-SUMO vector by restriction-free cloning^30^ for expression and purification in *E. coli* (Fig. 1c). Similarly to what observed for the full-length protein, CSB-HD incubation with 5’-tailed rDNA and its complementary strand did not fully resolve the G4 structure, but only promoted the G4 resolution and consequent double strand formation of about 15% compared to the control without incubation with CSB-HD (Fig. 1f, Supplementary Fig. S4a, Supplementary Table S3).

Moreover, negligible G4-resolvase activity on both 5’-tailed (Supplementary Fig. S4b, c, Supplementary Table S3) or untailed c-KIT1 G4 (Supplementary Fig. S4d, e, Supplementary Table S3) was also observed in the presence of CSB-HD, suggesting that CSB DNA-binding domain retains similar activity as the full-length protein. Overall, we confirmed that CSB is not a good G4-unwinder with only weak activity in promoting the resolution of tailed-rDNA G4s and that such activity can be recapitulated by the DNA binding domain of the protein alone.

### CD analysis fails to identify structural features that are peculiar of rDNA G4s

Since CSB G4-resolvase activity towards rDNA G4 revealed no dependency on incubation times or CSB concentrations (Supplementary Fig. S5a, b), we hypothesised that CSB might only recognise and resolve a subpopulation of G4-structures formed exclusively within rDNA under our experimental conditions. To investigate this, we performed circular dichroism (CD) analysis of the different rDNA sequences tested including and additional rDNA-3 G4-forming sequence^12^ (Supplementary Fig. S6 and Supplementary Table S1 for the sequences tested in this section). CD analysis revealed that rDNA G4s are mostly folded in a parallel conformation as assessed by the maxima ∼263 nm and a minimum ∼240 nm^31^ in the presence or absence of the tail 5’- or 3’-tail (Fig. 2a, Supplementary Fig. S6b, c, d, e, f) with exception of untailed rDNA which revealed a mixed topology^32^ (Supplementary Fig. S6a). The presence of a shoulder at ∼290 nm suggested a small population of rDNA G4s folded into an anti-parallel conformation (Fig. 2a, Supplementary Fig. S6b, c), which could be responsible of the helicase activity. However, the inability of CSB to resolve HRAS or hTELO sequences, which fold into anti-parallel or mixed type conformation respectively^33^, rules out a conformational based selectivity of CSB.

**Figure 2.**
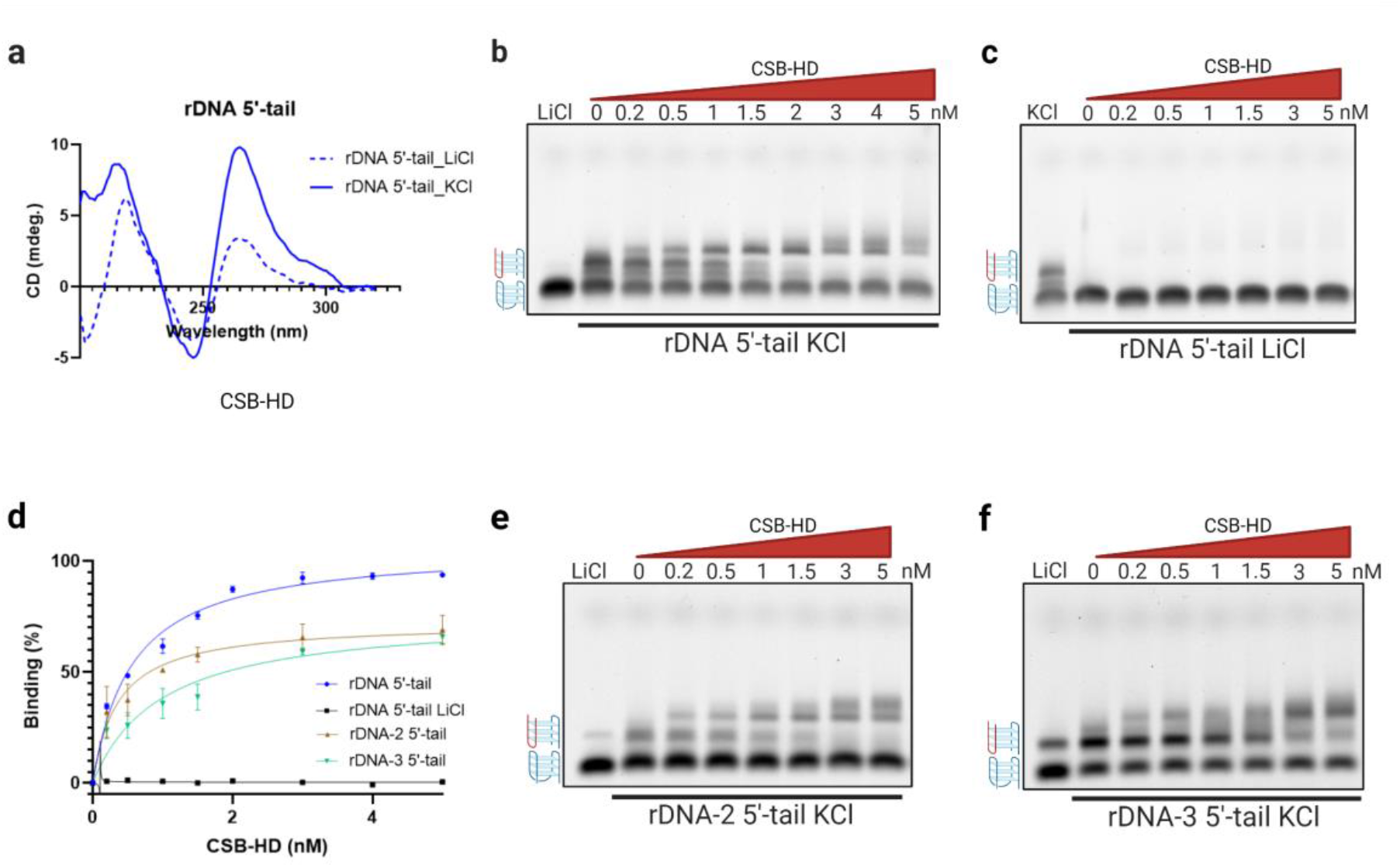
CSB selectively binds bimolecular rDNA G4s. (**a**) Circular dichroism (CD) analysis of rDNA 5’-tail sequence in LiCl or KCl buffer. The recorded spectra represent the average of three different reads after subtraction of the buffer signal. (**b**) EMSA gel on 5’-tailed rDNA G4 using 0 to 5 nM CSB-HD in KCl buffer. The first lane is a control with the oligonucleotide annealed in LiCl. (**c**) EMSA gel on 5’-tailed rDNA G4 using 0 to 5 nM CSB-HD in LiCl buffer. (**d**) Binding curves expressing the percentage of different oligonucleotides bound by increasing concentrations of CSB-HD (0 to 5 nM) in KCl or LiCl buffers. (**e**) EMSA gel on 5’-tailed rDNA-2 G4 using 0 to 5 nM CSB-HD in KCl buffer. The first lane is a control with the oligonucleotide annealed in LiCl. (**f**) EMSA gel on 5’-tailed rDNA-3 G4 using 0 to 5 nM in KCl buffer. The first lane is a control with the oligonucleotide annealed in LiCl. Unimolecular G4s are indicated as blue strand G4s while multimeric G4s are indicated with red and blue strand G4s. All the experiments were performed in duplicates and the binding K_ds_ were calculated based on two separate EMSAs experiments using one site specific binding equation using GraphPad Prism 9.0.1 software and 95% CI.

### CSB selectively binds intermolecular rDNA G4s with picomolar affinity

Given that a modest helicase activity limited to rDNA G4s has been observed in our helicase assays, we decided to further investigate the interaction between rDNA G4s and CSB by measuring its binding affinity towards the same panel of rDNA and non-rDNA G4s tested in the helicase assay by Electromobility-Shift Assay (EMSA)^34^. Interestingly, with the agarose gel used for this experiment, we were able to distinguish multiple bands in the free rDNA G4 sample in KCl buffer that were not detected when the oligonucleotide was annealed in LiCl (Fig. 2b, c, Supplementary Fig. S7b, c, d, e). Specifically, we observed two slow moving bands with apparent molecular weight of either twice or four times the one expected by the free oligonucleotide (Supplementary Fig. S7a). Based on this, we hypothesised that these slow-moving bands could be indicative of intermolecular G4s (bi- and tetra-molecular), whilst the more intense fast-moving band detected also in LiCl was to be ascribed to the unimolecular G4 or single stranded DNA. We then incubated this rDNA sequence annealed in KCl (see Supplementary Table S1 for all the sequences used in this section) with increasing concentrations (0-5 nM) of CSB-HD (Fig. 2b). We observed that CSB interacted selectively with the intermolecular G4 bands, displaying an astonishing picomolar affinity for multimeric rDNA G4s with a measured K_d_ of 557.5 pM [442.2 - 698.0 pM – 95% CI] (Fig. 2b, d), whilst displaying negligible binding against the fast-moving intramolecular G4 band. To further confirm the selective affinity for multimeric rDNA G4s, we also performed the same experiment under LiCl conditions, where no multimeric G4s bands are detected, with negligible binding observed under these conditions (Fig. 2c, d), suggesting high specificity for intermolecular G4s over intramolecular ones.

To further validate such high specificity towards intermolecular G4s, we then tested the binding of CSB against different 5’-tailed rDNA G4-forming sequences. To do so, we tested 5’-tailed rDNA-2 and 5’-tailed rDNA-3^12^ (see Supplementary Table S1 for the sequences used in this section). These rDNA sequences also displayed formation of intermolecular G4s under K^+^ stabilisation conditions, which are bound by CSB with K_d_s of 359.9 pM [205.2 - 594.1 pM – 95% CI] (Fig. 2d, e) and 977.1 pM [476.3 - 1924 pM – 95% CI], respectively (Fig. 2d, f).

We next sought to explore whether CSB is able to bind to intramolecular G4 structures. To achieve this, we tested CSB-HD binding against 5’-tailed c-KIT1, c-MYC, HRAS and hTELO (Supplementary Fig. S8a, b, c, d) and interestingly, we could not detect any interaction between CSB-HD and these G4s under the same conditions used to detect the interaction with intermolecular rDNA G4s. Conversely, we could observe for all the sequences tested formation of complexes with CSB at concentration between 50 and 200 nM, which suggested that at higher concentration of CSB interaction with intramolecular G4s can be observed. We also tested the ability of CSB-HD to interact with a non-G4-forming single stranded (ss) DNA sequence (see Supplementary Table S1 for the sequences used in this section) and, similarly to what observed for intramolecular G4s, the binding shifts are detectable only at CSB-HD concentration of 50 nM or higher (Supplementary Fig. S8e), suggesting that the interaction with intramolecular G4s is not specific over ssDNA and is ∼3 orders of magnitude lower than what observed for intermolecular G4s. Similar binding of CSB-FL (Supplementary Fig. S7b, c) or CSB-HD (Supplementary Fig. S7d, e) to intramolecular G4s at protein concentrations > 50nM could also be detected for rDNA sequences both in KCl or LiCl buffers, further demonstrating that this interaction is unable to discriminate between ssDNA and intramolecular G4s.

### CSB binds intermolecular G4s with a modest preference for a 3’ orientation

We next wanted to investigate if the positioning of the tail influences the binding of CSB on rDNA G4s and if the presence of the tail is necessary such interaction to occur. To do this, we tested the binding of CSB-HD on 3’-tailed rDNA and 3’-tailed c-MYC sequences, observing picomolar affinity on 3’-tailed rDNA with K_d_ of 175.5 pM [82.71 - 315.9 pM – 95% CI] (Fig. 3b, c) and negligible binding to 3’-tailed c-MYC (Supplementary Fig. S9a). The affinity of CSB for 3’-tailed rDNA is ∼3 folds higher than for its 5’-tailed counterpart, indicating that CSB may have a preference for intermolecular G4s resolvase activity with a 3′-to 5′-polarity. To further investigate if the tail was essential for such interaction, we also tested untailed rDNA, rDNA-2 and c-MYC sequences. Interestingly, we observed negligible formation of the intermolecular G4 structures with consequent no binding of CSB-HD on the three sequences (Fig. 3c, d, Supplementary Fig. S9b, c), suggesting that either a 5’ or a 3’-tail is necessary for the intermolecular G4s to form in the first place for these particular sequences under our experimental conditions. These results also suggested that the lack of rDNA G4 resolution observed in absence of the tail by helicase assays (Supplementary Table S2, Supplementary Figs. S1c, S2b) is most likely due to the lack of multimeric G4s formation of those sequences rather than to a strict requirement of a ssDNA tail to initiate the unwinding process as previously suggested^15^.

**Figure 3.**
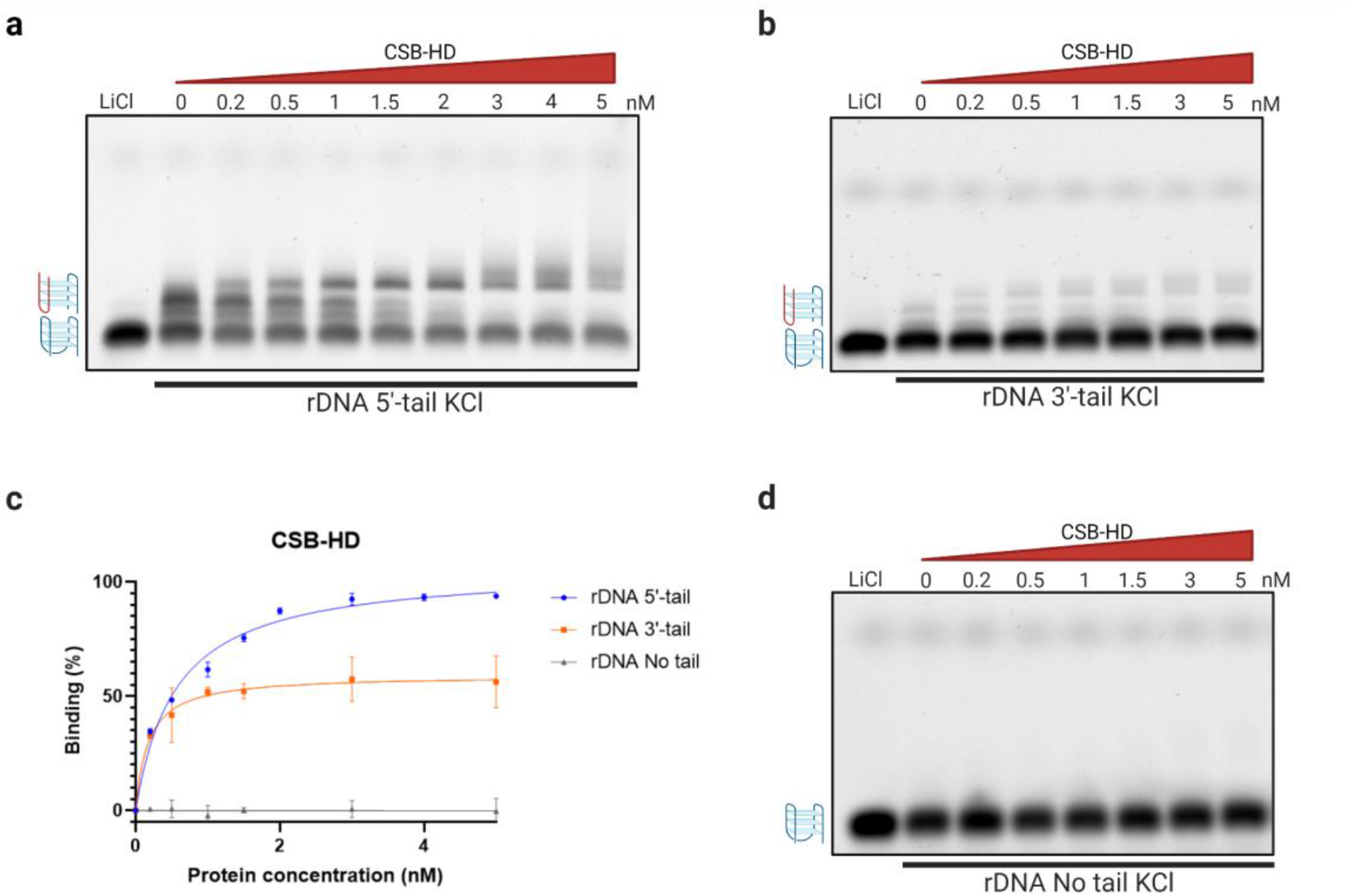
A ssDNA tail is essential for intermolecular rDNA G4s to form. EMSA agarose showing the formation of intermolecular rDNA G4 in KCl buffer in presence of a 3’-tail. Unimolecular G4s are indicated with blue strand G4s while intermolecular G4s are indicated with red and blue strand G4s. (**a**) EMSA with 5’-tail rDNA. (**b**) EMSA with 3’-tail rDNA. (**c**) Binding curves expressing the percentage of oligonucleotide bound by increasing concentrations of CSB-HD (0 to 5 nM) in KCl buffer. All the experiments were done in duplicates and plotted using one-site specific binding equation. (**d**) EMSA with untailed rDNA.

Moreover, from the gels we measured that the intensity of the bands related to the more stable unimolecular G4s are between 2 to 8 times higher than the multimeric G4 bands, suggesting that only small amount of intermolecular G4s is folded under these conditions. Therefore, the modest (∼15%) rDNA G4-resolvase activity observed after incubation with CSB (Fig. 2e, f, Supplementary Tables S2, S3) is likely to represent the specific resolution of the intermolecular G4s that are present in the solution in a smaller proportion. Taken together these results showed a surprisingly selective and strong interaction of CSB with bimolecular and tetramolecular rDNA G4s, which might be important to regulate rDNA G4 homeostasis and to prevent the premature ageing phenotype in CS patients lacking functional CSB.

### CSB binds and resolves intermolecular G4s

Agarose gel revealed the formation of multiple bands for all the tailed rDNA G4 sequences tested under KCl conditions (Figs. 2, 3), which we ascribed to multimeric (mainly bimolecular, Supplementary Fig. S7a) G4 structures and suggested a selective binding of CSB to multimeric G4s. To further demonstrate this, we tested the binding affinity of CSB-HD for a non-rDNA bimolecular G4 structure. Specifically, we used a 5’-tailed (or untailed) *O. nova* d(G4T4G4)^35,36^ telomeric sequence in our EMSA assay under the same conditions described for rDNA. Similar to what observed for 5’-tailed rDNAs, the 5’-tailed d(G4T4G4) sequence formed a slower moving band when annealed in KCl but not in LiCl (Fig. 4a), which we ascribed to the bimolecular G4s formed upon K^+^ stabilisation. CSB-HD was able to selectively interact with the dimeric G4s formed by the tailed d(G4T4G4) with an even stronger affinity than what observed for rDNA, yielding a measured K_d_ of 9.878 pM [2.872 - 23.91 pM – 95% CI] (Fig. 7a, b). Similarly, for the untailed d(G4T4G4) we could not detect formation of the slower moving band either in KCl or LiCl buffer (Fig. 4c), indicating that the bimolecular G4 was not formed in the absence of the tail under our experimental conditions. Because of the *O. nova* telomeric sequence presents only two guanine tracts, unimolecular G4s are unable to form within this sequence, indicating that the bands that fast-moving observed on the gel were to be ascribed to ssDNA. Accordingly, no interaction was observed between CSB-HD and the untailed d(G4T4G4) (Fig. 4b, c), confirming our hypothesis on the ability of CSB to interact selectively with intermolecular G4s.

**Figure 4.**
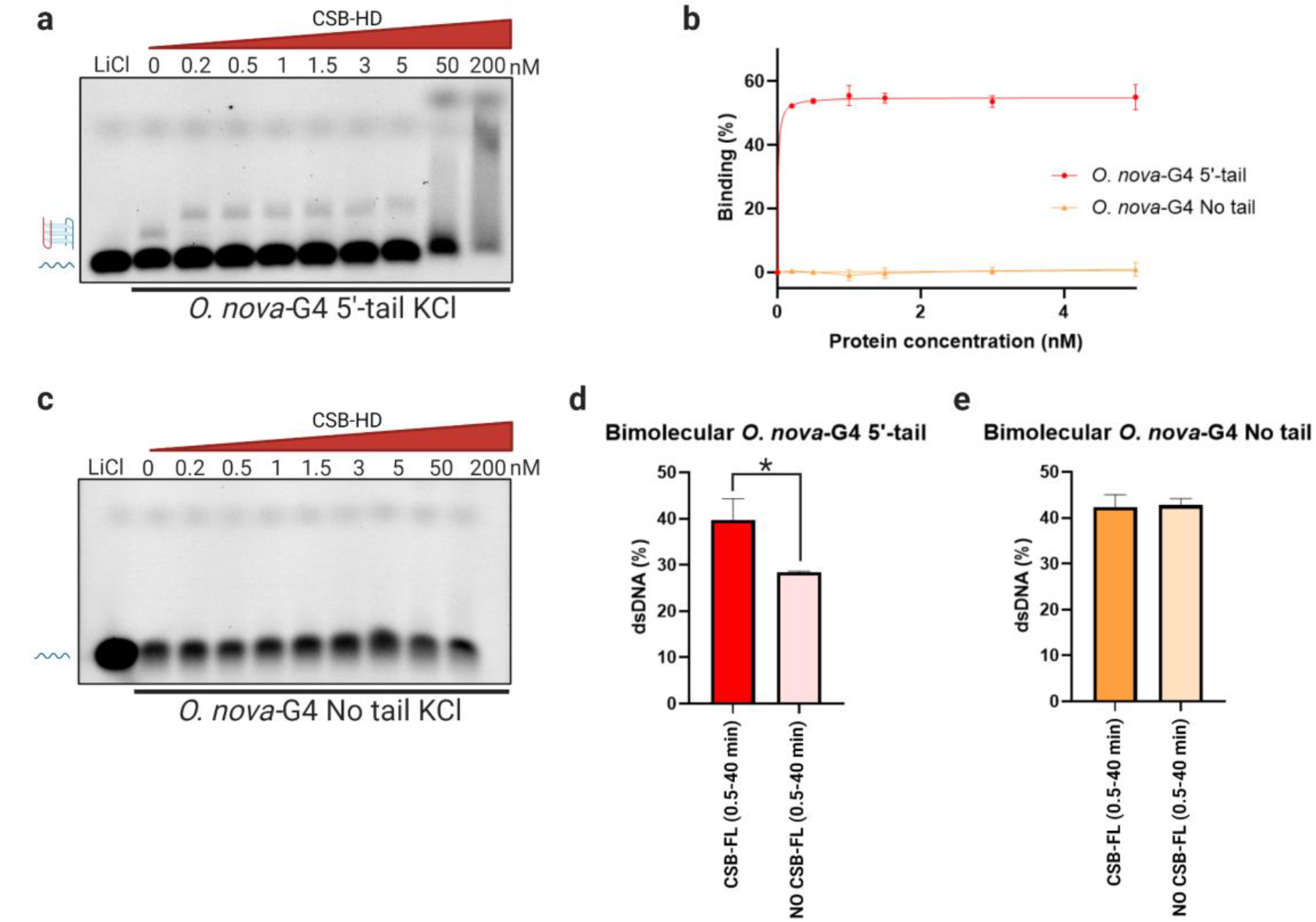
CSB selectively binds and resolves bimolecular G4s. (**a**) EMSA using 5’-tail *O. nova* bimolecular-G4 in KCl buffer. ssDNA bands are indicated with a blue linear strand while bimolecular G4s are indicated with red and blue strand G4. (**b**) Binding curves expressing the percentage of oligonucleotide bound by increasing concentrations of CSB-HD (0 to 5 nM) in KCl buffer. EMSAs were repeated in duplicates and plotted using one-site specific binding equation. (**c**) EMSA using untailed *O. nova* bimolecular-G4 in KCl buffer. The ssDNA bands are indicated with a blue linear strand. (**d**) Column graph of quantified helicase assay gel after incubation of 5’-tail *O. nova* bimolecular-G4 with or without CSB-FL in KCl buffer. (**e**) Column graph of quantified helicase assay of untailed *O. nova* bimolecular-G4 with or without CSB-FL in KCl buffer. The results are expressed as percentage of dsDNA formation. All quantified helicase assays were based on the average of three independent experiments ± SEM. Significance was calculated based on two-tailed Student’s t-test. Asterisks indicate statistical significance at 95% CI between the data with *p < 0.05.

To corroborate that CSB was able to selectively resolve intermolecular G4s we performed gel-based helicase assay on both 5’-tailed and untailed d(G4T4G4) sequence. As expected, for 5’-tailed d(G4T4G4) we observed 11.4% acceleration in dsDNA formation after incubation with CSB-FL compared to the control without CSB (Fig. 4d, Supplementary Fig. S10a), which is in agreement with the small fraction of bimolecular G4 formed under our experimental conditions. Furthermore, negligible acceleration in dsDNA formation was detected for the untailed d(G4T4G4) substrate (Fig. 4e, Supplementary Fig. S10b), confirming that in the absence of the bimolecular G4 substrate no helicase activity is elicited by CSB. Overall, these data demonstrated that CSB interacts selectively, with strong picomolar affinity, with the bimolecular G4 formed by 5’-tail *O. nova* d(G4T4G4) telomeric sequence and that this interaction can promote the selective resolution of such intermolecular G4 in ATP-independent fashion when in the presence of its complementary DNA strand.

### Treatment with PDS or CX-5461 displaces CSB-rDNA G4 interaction

Previous studies revealed that treatment with G4-stabilizing molecules, such as Pyridostatin (PDS) and CX-5461 promotes accelerated aging through rDNA G4 stabilisation in human cells and *C. elegans* models^15^. Therefore, we asked whether these G4-ligands can displace CSB bound to its intermolecular rDNA G4s substrate, which could explain the premature ageing phenotype observed in these models upon PDS or CX-5461 treatment^15^ as well as CSB displacement from the nucleolus observed by confocal imaging using CSB-GFP upon treatment with CX-5461^6^. To test this, we investigated if PDS^18^ and CX-5461^19^ were able to displace the CSB-rDNA G4 interaction in a dose dependent fashion. We performed a competitive EMSA where we incubated 2 nM CSB-HD pre-bound to rDNA G4 with increasing concentrations of either PDS or CX-5461 (0-5000 nM). As displayed in Figure 5a and 5b, we were able to observe replenishment of the free intermolecular rDNA G4s bands in a dose dependent fashion (Fig. 5a, b), demonstrating that G4s ligands can compete with CSB for binding to intermolecular rDNA G4s under our experimental conditions. CX-5461 has often been considered a transcriptional inhibitor in previous reports^6,15,37,38^, albeit it has been previously designed and recently further validated as an effective G4-ligand^19^. Our results demonstrate that CX-5461 can outcompete CSB for binding to intermolecular rDNA G4s, suggesting that both the premature ageing phenotype and the CSB nucleolus displacement observed upon treatment with this ligand are more likely due to its G4-binding ability rather than transcriptional inhibition, similarly to what observed with PDS.

**Figure 5.**
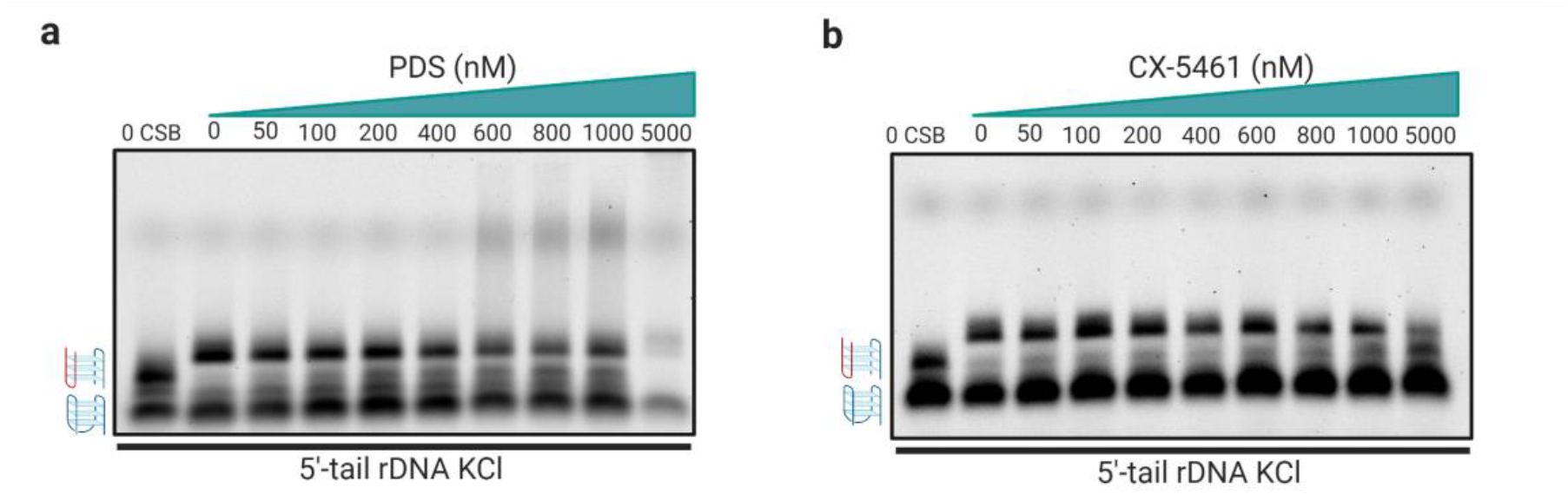
The interaction between CSB and bimolecular G4 is displaced by G4-ligands. (**a**) EMSA agarose gel obtained after incubation of 5’-tail rDNA G4 with 2 nM CSB-HD in presence of increasing concentrations of PDS (0 to 5000 nM) in KCl buffer. The first lane is a control of free 5’-tail rDNA showing the formation of unimolecular G4 (blue strand G4) and multimeric G4 (G4 formed within the red and blue strands). (**b**) EMSA agarose gel obtained after incubation of 5’-tail rDNA G4 with 2 nM CSB-HD in presence of increasing concentrations of CX-5461 (0 to 5000 nM) in KCl buffer. The first lane is a control of free 5’-tail rDNA showing the formation of unimolecular G4 (blue strand G4) and multimeric G4 (G4 formed within the red and blue strands).

## Discussion

As identified by high-throughput sequencing mapping of G4s (G4-seq), these structures can be widely distributed throughout the human genome^39^. Nevertheless, RNA-seq experiments in cells that lack functional CSB demonstrated a specific increase of transcriptional stalling specifically at rDNA G4s, without significant transcriptional alteration at other G4-forming sites of the genome^15^. This hinted at a close link between CSB and rDNA G4 that upon loss of function of this protein triggers transcriptional stalling selectively at rDNA G4s, eliciting a series of biological responses that are typical of ageing^15,16,17^. Previous studies suggested that CSB can act as an ATP-independent G4-helicase and proposed that lack of CSB G4-resolvase activity was responsible for this phenotype^15^. However, CSB belongs to a family of chromatin remodelling protein and although the presence of a DEGH box in his helicase-like domain^23^ (519-1002 aa, Fig 1a), CSB has not been classified as a canonical helicase^40^. Furthermore, previous reports showed that CSB was a modest G4-resolvase, which is in contrast to the strong phenotype observed in Cockayne Syndrome patients. Therefore, we decided to further investigate the nature of the interaction between CSB and rDNA G4s. Our helicase assays revealed that the apparently modest G4-resolution ability of CSB is strictly specific to rDNA G4s (Fig. 1e, f).

Although rDNA selective, the modest CSB ability to resolve rDNA G4s was still incongruous with the strong G4-mediated transcriptional stalling observed at rDNA in CSB impaired cells, which prompted us to further investigate the ability of CSB to bind G4 structures by EMSA. The agarose gel used for this assay allowed us to identify alternative conformations for all the rDNA G4-forming sequences tested either 5’- or 3’-tailed. Since these alternative conformations were only visible in KCl stabilising buffer as slow-moving bands on a gel (Figs. 2b, e, f, 3b, Supplementary Fig. S7), we reasoned that rDNA sequences might fold into bimolecular and tetramolecular G4s (Supplementary Fig. S7a). Surprisingly, we observed an astonishing picomolar affinity of CSB for intermolecular G4s (Figs. 2b, d, e, f, 3b, c) whilst no binding was detected against the intramolecular G4s formed by the very same sequences. This suggested that intermolecular G4 structures might be formed in cells within the nucleolus and that CSB can act as a dedicated protein to both bind and resolve these structures in living cells.

This hypothesis was further confirmed by the analysis of CSB binding towards a non-rDNA bimolecular G4 formed by the *O. nova* d(G4T4G4)^35,36^ sequence. We observed that CSB binds and resolves also this bimolecular G4 with an even stronger affinity (∼10 pM, Fig. 4a, b, d), corroborating the selective and high binding affinity of CSB to intermolecular G4s.

Despite the ability of G-rich sequences to form intermolecular G4s has been known for decades, G4-related biological studies have focused almost exclusively on intramolecular G4s. This assumption has often been justified by the low probability of intermolecular G4 to form within genomic, chromatinised DNA. However, our study demonstrates that CSB has an astonishing affinity for intermolecular G4s with a ∼1000-fold higher selectivity over intramolecular G4s or ssDNA, suggesting that rDNA sequences are likely to adopt multimeric G4-conformations in cells as well as they do *in vitro* and act as a natural substrate of CSB. Our findings, in combination with previous reports that have demonstrated transcriptional stalling at rDNA sites in CSB impaired cells, hint at a specific biological role that CSB has in the maintenance on rDNA transcription homeostasis through a selective interaction with intermolecular G4s.

Previous experiments using *C. elegans* revealed that treatment with G4-stabilising ligands such as PDS^18^ and CX-5461^19^ accelerates the premature ageing phenotype in this model^15^, whilst confocal imaging of human CS1AN cells expressing CSB-GFP revealed that CX-5461 can displace CSB from the nucleolus^6^. These observations in combination with our data demonstrating that G4-ligands can outcompete CSB for binding to intermolecular G4s (Fig. 5), support a model where the interaction between CSB and intermolecular rDNA G4s is somewhat essential to prevent premature ageing development (Fig. 6).

**Figure 6.**
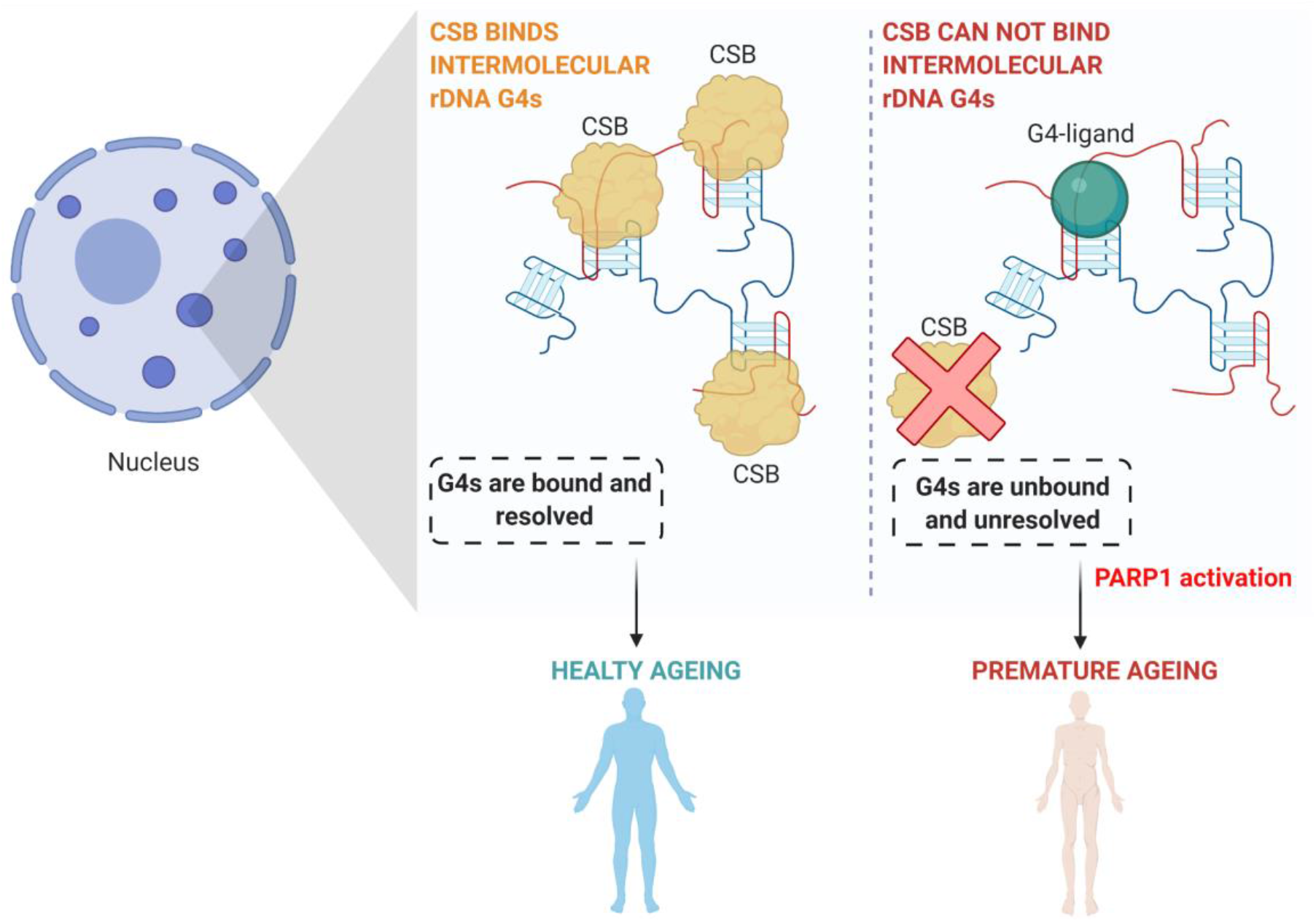
CSB-inter-molecular rDNA G4 interaction is essential to prevent premature ageing development. Model showing the formation of intermolecular G4 within the condensed nucleoli (blue dots in the nucleus). (**a**) On the left, the zoomed nucleoli present active CSB that binds selectively rDNA intermolecular G4s (formed within the red and blue strands) with strong picomolar affinity and promotes their resolution. Controlled G4-handeling given by CSB promotes healthy ageing. (**b**) On the right, when CSB is not functional or the interaction has been displaced by G4-ligands, rDNA intermolecular G4s remain unbound by CSB and unresolved. They are recognised as DNA damage, activating PARP1 and consequent cellular stress typical of the premature ageing phenotype.

In summary, we presented the first report of an endogenous protein able to bind with high affinity and specificity intermolecular G4 structures over intramolecular ones. Nucleoli are dense nuclear regions that accommodate the highly G-rich rDNA, creating the ideal environment to promote formation of intermolecular G4 structures. Our data on the selectivity and the high affinity of CSB for intermolecular G4s suggests that these structures are likely formed in the nucleolus of human cells and that CSB is a dedicated protein to regulate their prevalence and functions, which is particularly relevant in the context of premature ageing.

## Methods

### CSB-FL expression and purification

The construct pFASTBac-HA-CSB-His6 encoding the full length CSB protein (1-1493 aa) with a N-terminal hemagglutinin antigen (HA) epitope and a C-terminal histidine (His6) tag for overexpression in baculovirus system was kindly donated by Scheibye-Knudsen group. The CSB-FL protein was expressed in *Spodoptera frugiperda (Sf9)* system by adapting previously reported protocols^21,22^. Briefly, suspension cultures of *Sf9* insect cells at 8×10^5^ cells/ml were infected for 72 hours with the recombinant baculovirus. At 3 days post-infection, cells were collected and harvested by centrifugation at 4°C for 10 min at 1500xg. The pellet from 2 litres cells was washed twice with ice-cold phosphate-buffered saline before resuspending the cells into 8 (packed cells) volumes of ice-cold lysis buffer and protease inhibitors (25 mM Tris-HCl pH 9.0, 1 mM EDTA, 10% glycerol, 1% Triton-x, 300 mM KCl, 1 mM TCEP, 0.1 mM PMSF, 0.2 μM chemostatin, lupeptin, antipapain and pepstatin A) supplemented with 5 mM MgCl_2_ and 5 KU Benzonase nuclease (Merck Millipore). After incubation at 4°C for 30 minutes to assure the complete lysis, cells were gently sonicated (sonicator from Thermo Fisher Scientific) in ice for 15 minutes, 20% amplitude, 2 seconds on, 58 seconds off. The suspension was then centrifugated at 4 °C for 40 minutes at 16000xg for 40 minutes to clarify the lysate. The supernatant was then incubated with 8 ml of HisPur Nickel-NTA Resin (Thermo Fisher Scientific) for the affinity purification. The bounded protein was washed 5 times with buffer containing 20 mM imidazole before elution in a dedicated buffer containing 25 mM HEPES pH 7.9, 10% glycerol, 0.01% Triton-x, 300 mM KCl, 250 mM imidazole, 1 mM TCEP and 0.1 mM PMSF. The eluate was directly loaded on 5 ml HiTrap Heparin High Performance column (Cytiva life sciences) equilibrated with heparin buffer (25 mM HEPES pH 7.9, 10% glycerol, 0.01% Triton-x, 300 mM KCl, 1 mM TCEP, 0.1 mM PMSF). The protein was then diluted using a linear gradient (0-100%) of heparin buffer containing 800 mM KCl. In the final step of purification, the heparin fractions were further purified using Superose 6 Increase 10/300 GL column (Cytiva life sciences) and eluted in gel filtration buffer (25 mM HEPES pH 7.9, 10% glycerol, 0.01% Triton-x, 200 mM KCl, 1 mM TCEP, 0.1 mM PMSF). The protein was then concentrated to 1 mg/ml using Amicon device 100K (Merck Millipore) and stored at -80°C. The stability of the recombinant protein during the purification was monitored by SDS-PAGE and Western blot analysis (WB) using a rabbit anti-CSB primary antibody (ABCAM: ab96089) and a goat anti-rabbit HRP secondary antibody (ABCAM: ab205718).

### CSB-HD cloning

To understand which was the most conserved region that recapitulates the CSB helicase-like domain in humans, we performed a multiple sequence alignment of different CSB homologues in different organisms, including humans, using Clustal Omega, EMBL-EBI^27,28^ software. Then, we performed a secondary structure prediction of the entire human CSB sequence (UniProtKB: Q03468) using PSIPRED 4.0 bioinformatic tool^29^. The most stable conserved gene sequence (CSB-HD: 498-1002 aa) was codon-optimised for bacterial expression and synthetised using Genscript service. The CSB-HD gene was then cloned into a pCS46_6xHistidine-SUMO vector for *E. coli* expression (this empty vector was kindly donated by Christian Speck group) by restriction-free cloning^30^ using forward primer: 5’-GTTCCAGCAGCAGACGGGAGGTAAAGTGCCGGGTTTCCTG-3’ and reverse primer: 5’-CAGCGGTGGCAGCAGCCAACTCTTAATACAGGTCATTGCTCTTAAAG-3’. Forward and reverse primers for the PCR reaction were designed using the online rf-cloning.org service and purchased from Eurogentec.

### CSB-HD expression and purification

The pCS46_6xHistidine-SUMO-CSB-HD plasmid was transformed into BL21(DE3) Competent *E. coli* cells (New England BioLabs) following the supplier heat shock protocol. The bacteria cells were then expanded to 4 litres in 2XYT media (Sigma Aldrich) at 37 °C and the expression of the protein was induced with 0.05 mM IPTG at 18 °C overnight. Cells were lysed in ice-cold lysis buffer (same buffer we used for the CSB-FL) supplemented with EDTA-free protease inhibitor tablets (Sigma-Aldrich), 5 mM MgCl_2_ and 5 KU Benzonase nuclease (Merck Millipore) for 30 mins at 4 °C. Cells were sonicated (sonicator from Thermo Fisher Scientific) in ice for 15 minutes, 20% amplitude, 5 seconds on and 55 seconds off. Clarified lysed was subjected to the same three-step purifications we used for the CSB-FL using the same buffers. The only difference was the using of a Superdex 200 Increase 10/300 GL column for the last size exclusion purification step (Cytiva life sciences). The protein was then concentrated to 1 mg/ml using Amicon device 30K (Merck Millipore) and stored at -80°C.

### Gel-based helicase assay

For helicase assay, the 5’-labelled oligonucleotides (see Table Supplementary S1) were annealed at a concentration of 100 nM in a buffer containing 25 mM HEPES pH 8.0, 100 mM KCl at 95 °C for 10 minutes, following slow cooling to room temperature overnight. The helicase reaction mixtures (20 μl) containing 1 nM of G4 and 5X fold excess of unlabelled complementary strand in helicase assay reaction buffer (20 mM HEPES pH 8.0, 40 ng/μl BSA, 1 mM DTT, 1 mM MgCl_2_, 25 mM KCl) were added with 10 nM of either CSB-FL and CSB-HD and incubated at 30 °C for different times (0.5-40 minutes). The reactions were stopped with 5 μl of stop solution containing 0.5% SDS, 50 mM EDTA and resolved in 10% polyacrylamide (19:1 acrylamide-bisacrylamide) gels at 100V for 1 hour in 0.5XTBE. The imaging of the gels was obtained using Typhoon FLA 9500 (GE Healtcare) set to detect Cy5 or FAM. The gel bands quantitation was processed using ImageJ software and analysed using GraphPad Prism 9.0.1.

### Circular dichroism (CD)

The annealing of the unlabelled oligonucleotides (see Table Supplementary S1) was performed in 10 mM Tris-HCl pH 7.4, 100 mM KCl or LiCl buffer at 95 °C for 10 minutes following slow cooling to room temperature overnight. The oligonucleotides were annealed at a final concentration of 10 μM. CD spectra were recorded on a JASCO J-810 circular dichroism spectrophotometer using a 1 mm path length quartz cuvette. CD measurements were performed at 298 K over a range of 200-320 nm using a response time of 2 s, 1nm pitch and 0.5 nm bandwidth. The recorded spectra represent the average of three different reads. The absorbance of the buffers was subtracted from the recorded spectra.

### Electromobility-shift assay (EMSA)

5’-(Cy5 or FAM)-fluorophore labelled oligonucleotides (see Table Supplementary S1) were annealed at a final concentration of 500 nM in 25 mM HEPES pH 8.0, 100 mM KCl (or 100 mM LiCl) buffer at 95 °C for 10 minutes following slow cooling to room temperature overnight. 10 μl reaction mixtures containing EMSA reaction buffer (20 mM HEPES pH 8.0, 40 ng/μl BSA, 1 mM DTT, 1 mM MgCl_2_, 100 mM KCl or LiCl), 25 nM of fluorophore labelled oligonucleotides and increasing concentrations of either CSB-FL or CSB-HD were incubated at 30 °C for 30 minutes. The reactions were stopped by addition of 2 μl of no SDS-purple gel loading dye (New England BioLabs) or 50% glycerol. The samples were then separated on a 0.6% agarose gel at 100V for 50 min in 1XTBE. For the competitive EMSAs, the reactions were performed using the same EMSA reaction buffer but with fixed concentration of CSB-HD (2 nM) and 5’-tailed rDNA G4 (25 nM). The samples were incubated at 30°C for 30 minutes before to add the desired concentration of Pyridostatin (PDS) or CX-5461 (0-5000 nM) diluted in milliQ water for 2 hours at 30 °C. The reactions were then stopped by 2 μl of no SDS-purple gel loading dye (New England BioLabs) and separated on 0.6% agarose gel at 100V for 50 min in 1XTBE. For. The gels were visualised by GE Typhoon imager set to detect Cy5 or FAM. The lane quantitation was processed using ImageJ software. The binding K_ds_ were calculated based on two separate EMSAs experiments using one site specific binding equation^41^ using GraphPad Prism 9.0.1 software and 95% CI.

### Agarose gel

To detect the formation of multimeric G4s we performed agarose gels. The unlabelled 5’-tail rDNA oligonucleotide (see Table Supplementary S1) was annealed at final concentration of concentration of 20 μM in buffers containing 25 mM HEPES pH 8.0 and 100mM LiCl or 25 mM HEPES pH 8.0 and either 100 mM KCl or 100 mM KCl + 30% PEG200 at 95 °C for 10 minutes, following slow cooling to room temperature overnight. Then 10 μM of each annealed oligonucleotide and 1μl of 50bp ladder (New England BioLabs) were loaded on a 0.6% agarose gel supplemented with 1X gel red and separated at 100V for 50 min in 1XTBE. The gels were visualised by G:BOX F3.

### Statistical analysis

The helicase assay data were presented as the mean ± SD in at least three separate experiments. Statistical significance was calculated based on two-way t-test using GraphPad Prism 9.0.1 software. The statistically significant difference was defined as P<0.05.

## Acknowledgment

This work and the M.D.A. group are supported by a BBSRC David Phillips Fellowship (BB/R011605/1/BB). D.L. is supported by an Imperial College DTP scholarship funded by the Chemistry Department. We thank Scheibye-Knudsen group (University of Copenhagen) for donating the plasmid for the expression of the CSB-FL in *S. frugiperda* (*Sf9*) and Evangelos Christodoulou, Philip Walker and Chloe Roustan (The Francis Crick Institute) for kindly helping with the expression of the CSB-FL during the closure of the Imperial College London’s Baculovirus Facility due the COVID-19 pandemic. We thank also Christian Speck group (Imperial College London) for generously providing the pCS46_6xHistidine-SUMO vector used to express the CSB-HD in *E. coli* and Vera Leber for the experimental support.

## Author contribution

M.D.A and D.L. conceptualized the work. D.L. performed and analyzed all the experiments. D.L. and M.D.A. interpreted the results and co-wrote the manuscript.

## Additional information

All the figures were created with BioRender.com.

