## Supplementary Information for "Cockayne Syndrome B protein selectively interacts and resolves intermolecular DNA G-quadruplex structures"

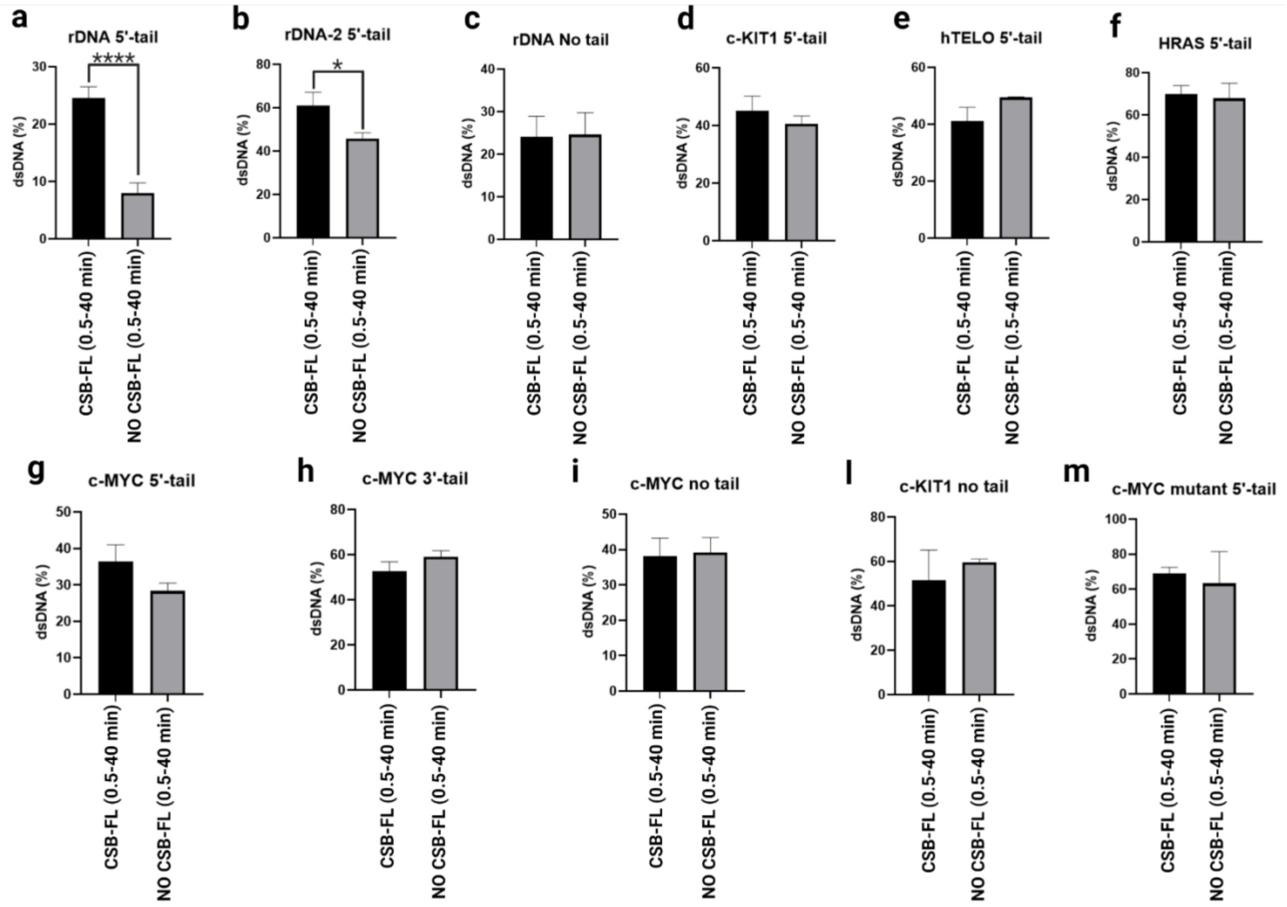


**Figure S1. CSB selectively resolves tailed rDNA G4s.** Column graph of quantified helicase assays with or without CSB-FL in KCl buffer. The results are expressed as percentage of dsDNA formation. (**a**) 5’-tail rDNA. (**b**) 5’-tail rDNA-2. (**c**) Untailed rDNA. (**d**) 5’-tail c-KIT1. (**e**) 5’-tail hTELO. (**f**) 5’-tail HRAS. (**g**) 5’-tail c-MYC. (**h**) 3-tail c-MYC. (**i**) Untailed c-MYC. (**l**) Untailed c-KIT1. (**m**) 5’-tail mut-c-MYC. All quantified helicase assays were based on the average of three independent experiments ± SEM. Significance was calculated based on two-tailed Student’s t-test. Asterisks indicate statistical significance at 95% CI between the data with ****p <0.0001 and *p < 0.05.

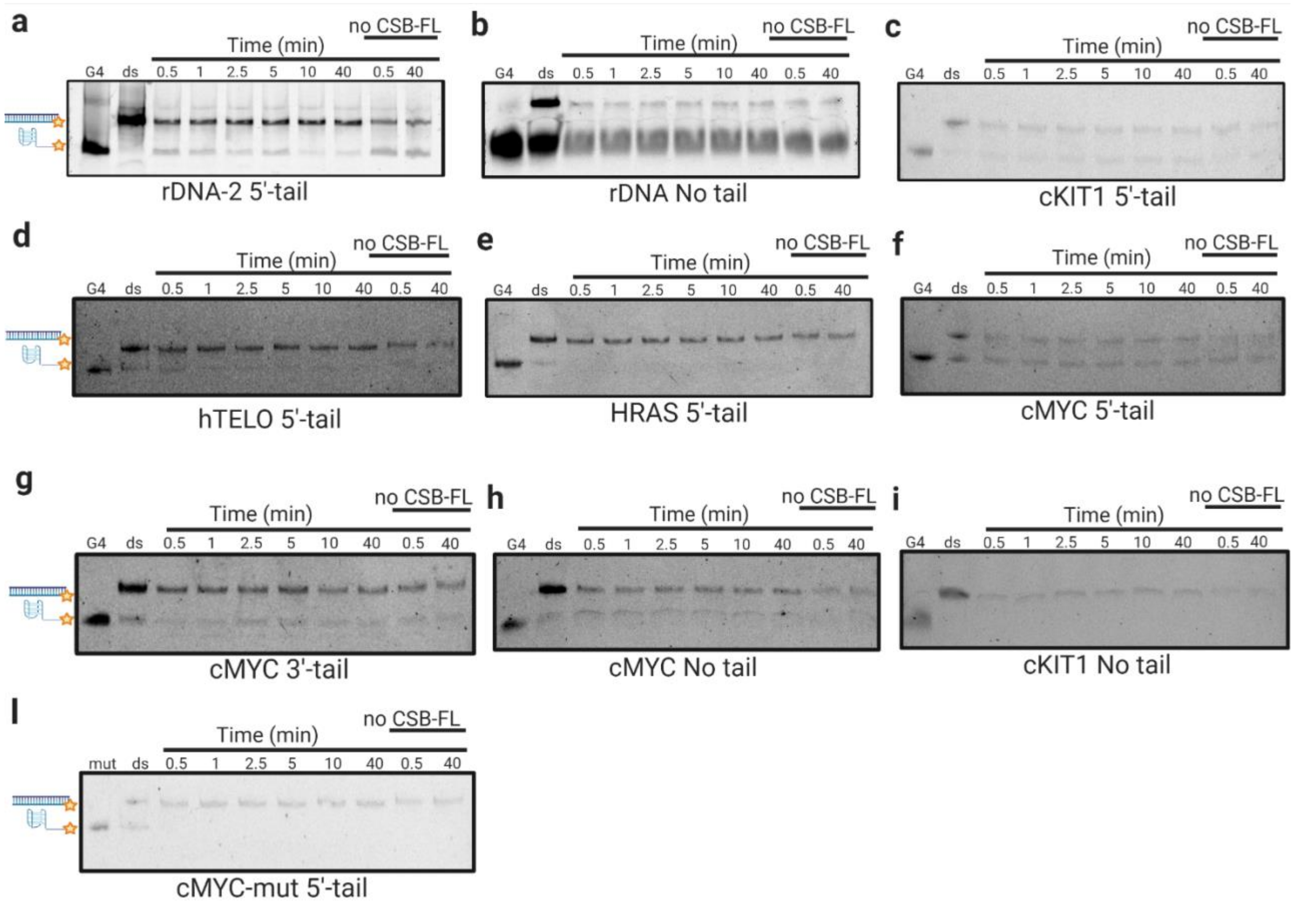


**Figure S2. Gel pictures of the helicase assays performed with CSB-FL.** (**a**) 5’-tail rDNA-2. (**b**) Untailed rDNA. (**c**) 5’-tail c-KIT1. (**d**) 5’-tail hTELO. (**e**) 5’-tail HRAS. (**f**) 5’-tail c-MYC. (**g**) 3-tail c-MYC. (**h**) Untailed c-MYC. (**i**) Untailed c-KIT1. (**l**) 5’-tail mut-c-MYC.

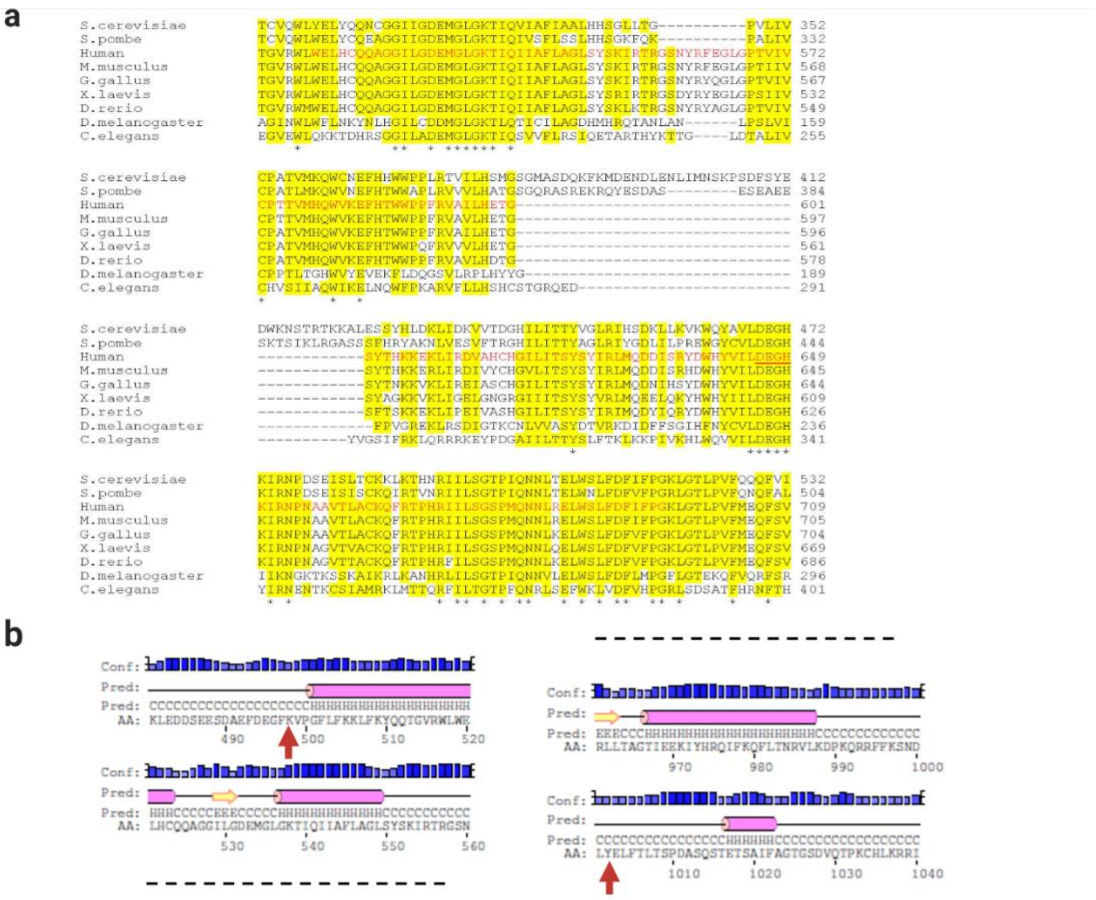


**Figure S3. CSB-HD isolation.** (**a**) Crop of the sequence alignment of the putative CSB helicase-like domain between different organisms (Clustal Omega, EMBL-EBI). In red is reported the *human* ATP-binding domain containing the DEGH box (646-649, underlined in red). * indicates 100% identity between the residues and in yellow are highlight the identical or similar residues. (**b**) Portion of the secondary structure prediction (PSIPRED 4.0) of the selected *human* helicase-like domain (498-1002 aa). The red arrows indicate the first N-terminal amino acid (K-498) and the last C-terminal residue (Y-1002). The α-helices are represented as pink cylinders while the β-coils are reported as yellow arrows.

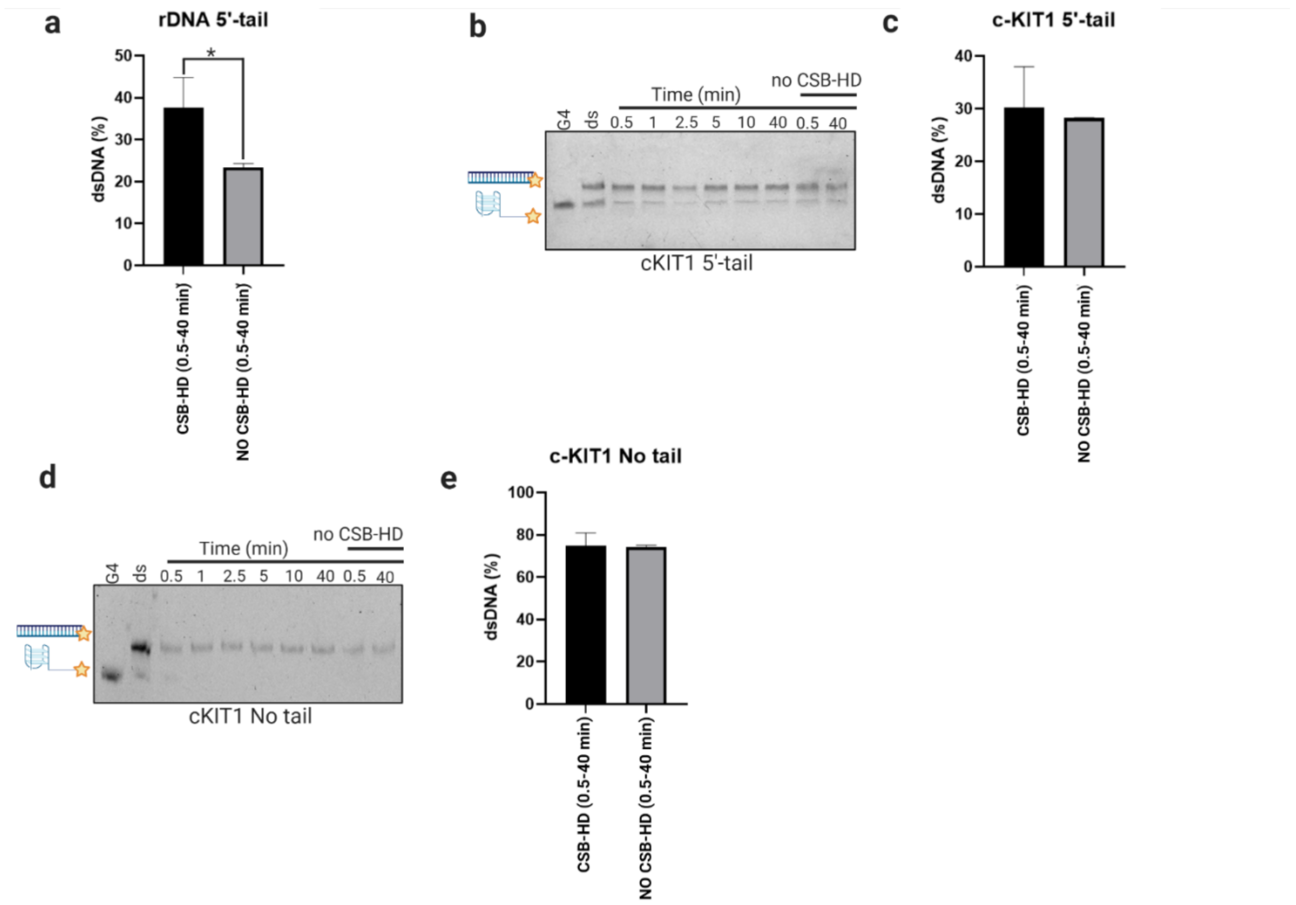


**Figure S4. CSB-HD retains the same activity as the FL protein in a G4-context.** (**a**) Column graph of quantified 5'-tail rDNA helicase assay with or without CSB-HD in KCl buffer. The results are expressed as percentage of dsDNA formation. (**b**) 5'-tail c-KIT1 helicase assay gel in presence or absence of CSB-HD (0.5-40 minutes). (**c**) Column graph of quantified 5'-tail c-KIT1 helicase assay. (**d**) Untailed c-KIT1 helicase assay gel in presence or absence of CSB-HD. (**e**) Column graph of quantified untailed c-KIT1 helicase assay. All quantified helicase assays were based on the average of three independent experiments ± SEM. Significance was calculated based on two-tailed Student's t-test. Asterisks indicate statistical significance at 95% CI between the data with $*p < 0.05$.

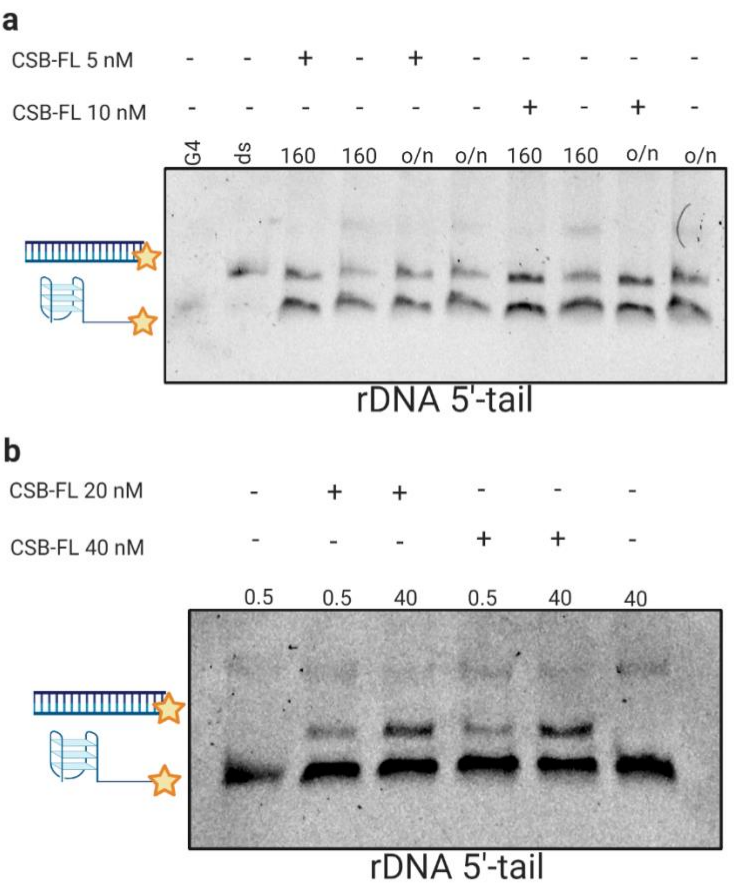


**Figure S5. CSB G4-resolvase activity towards 5'-tail rDNA G4s is not dependent on incubation times or CSB concentrations.** (**a**) Helicase assay gel in presence or absence of either 5 nM or 10 nM CSB-FL after 160 min or overnight (o/n) incubation time. (**b**) Helicase assay gel in presence of either 20 nM or 40 nM CSB-FL after 0.5 min or 40 min incubation time.

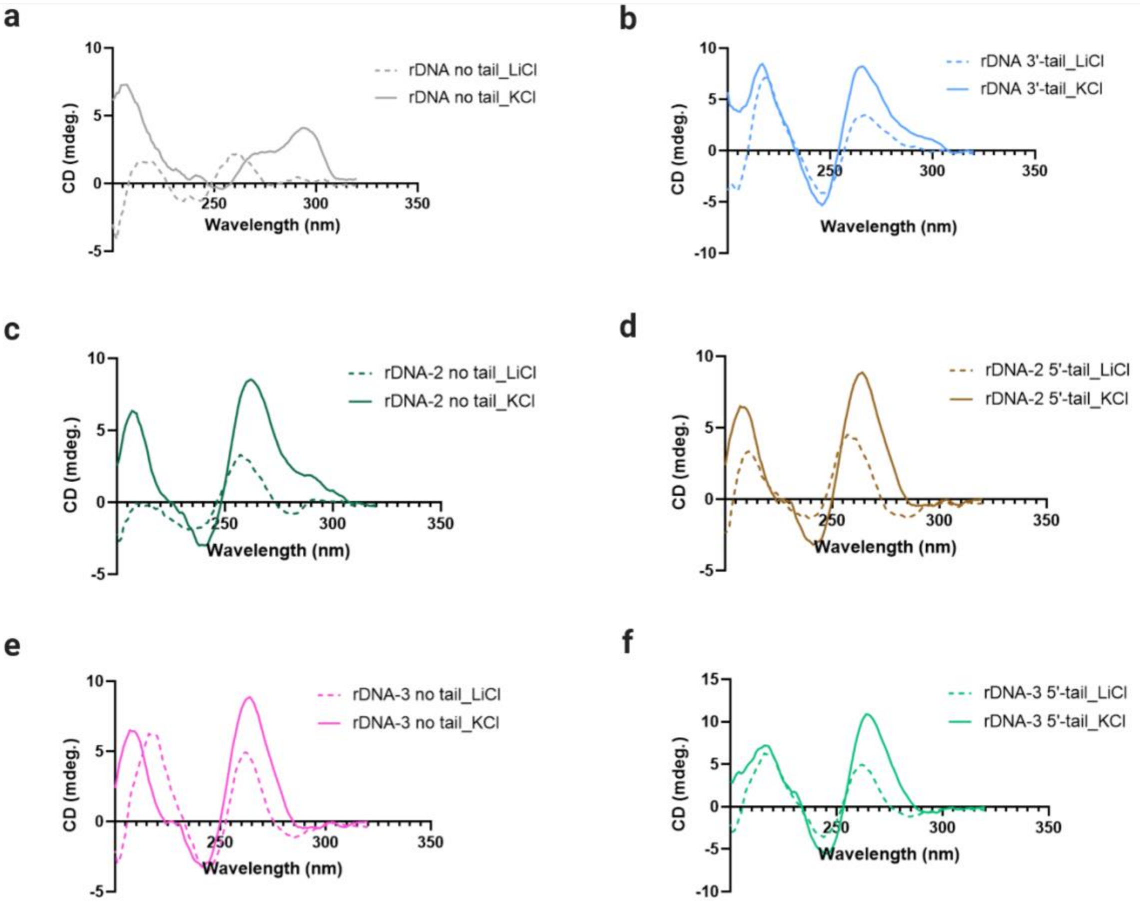


**Figure S6. Circular dichroism (CD) analysis of the different rDNA sequences in LiCl or KCl buffer.** (**a**) CD spectra rDNA No tail. (**b**) CD spectra rDNA 3'-tail. (**c**) CD spectra rDNA-2 No tail. (**d**) CD spectra rDNA-2 5'-tail. (**e**) CD spectra rDNA-3 No tail. (**f**) CD spectra rDNA-3 5'-tail. The recorded spectra represent the average of three different reads. The absorbance of the buffers was subtracted from the recorded spectra.

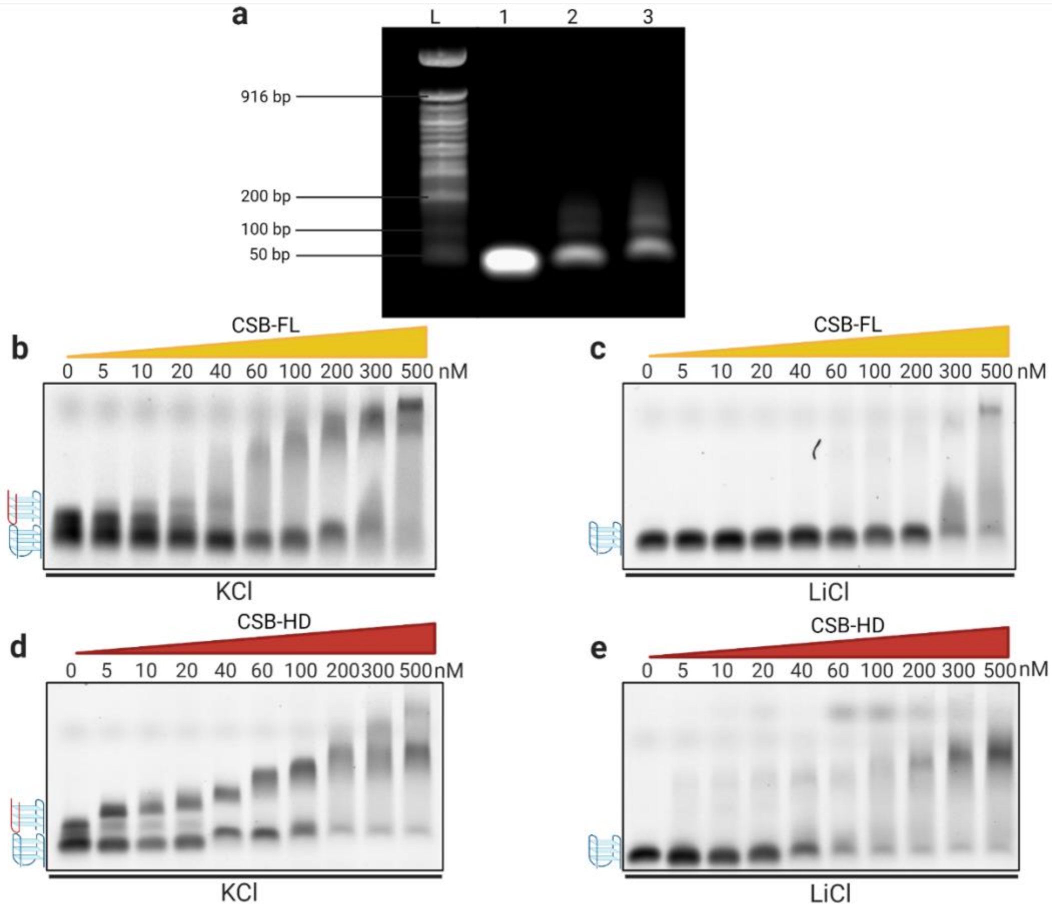


**Figure S7. EMSA Agarose revealed the formation of multimeric rDNA G4s formation.** (**a**) Agarose gel of unlabelled 5’-tail rDNA substrate annealed in buffer containing: 100mM LiCl (1), 100mM KCl (2) or 100nM KCl+30% PEG200 (3). L: ladder. (**b**) EMSA gel using high concentration (0 to 500 nM) CSB-FL in presence of 5’-tail rDNA in KCl buffer. (**c**) EMSA gel using high concentration (0 to 500 nM) CSB-FL in presence of 5’-tail rDNA in LiCl buffer. (**d**) EMSA gel using high concentration (0 to 500 nM) CSB-HD in presence of 5’-tail rDNA in KCl buffer. (**e**) EMSA gel using high concentration (0 to 500 nM) CSB-HD in presence of 5’-tail rDNA in LiCl buffer. Unimolecular G4s are indicated as blue strand G4s while multimeric G4s are indicated with red and blue strand G4s.

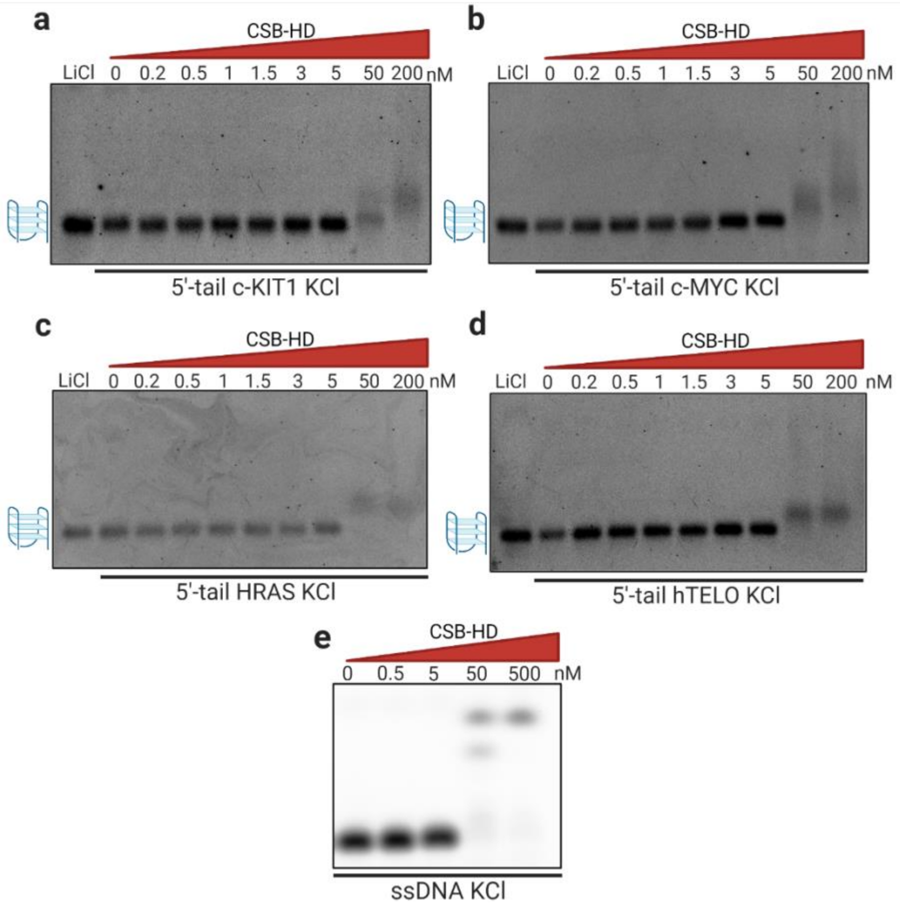


**Figure S8. CSB does not bind non-rDNA G4s at low concentration.** EMSA agarose incubating increasing concentration of CSB-HD (0 to 5 nM and 50-200 nM or 50-500 nM) with different non-rDNA G4 oligonucleotides under KCl conditions. (**a**) EMSA using 5’-tail c-KIT1. (**b**) EMSA using 5’-tail c-MYC. (**c**) EMSA using 5’-tail HRAS. (**d**) EMSA using 5’-tail hTELO. (**e**) EMSA using 5’-tail ssDNA as control. Unimolecular G4s are indicated as blue strand G4s.

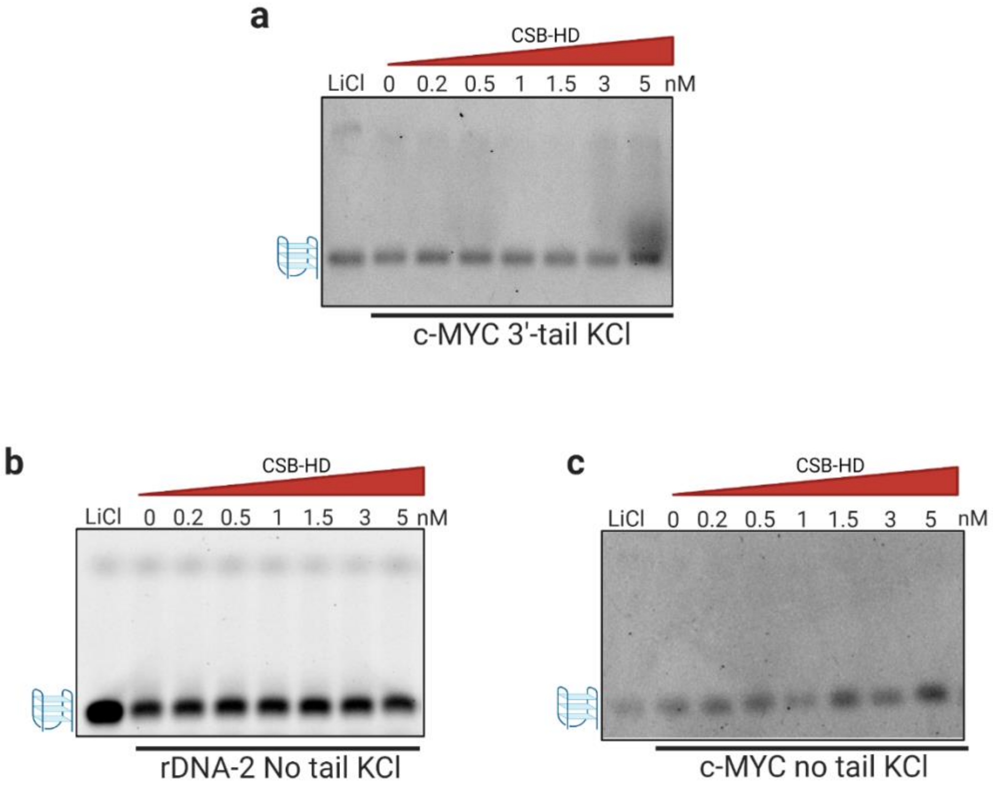


**Figure S9. Intermolecular G4s cannot form within rDNA sequences in absence of a tail.** EMSA agarose incubating increasing concentration of CSB-HD (0 to 5 nM) in presence of 3'-tailed intramolecular G4 or untailed oligonucleotides under KCl conditions. (**a**) EMSA using 3'-tail c-MYC. (**b**) EMSA using untailed rDNA-2. (**c**) EMSA using untailed c-MYC. Unimolecular G4s are indicated as blue strand G4s.

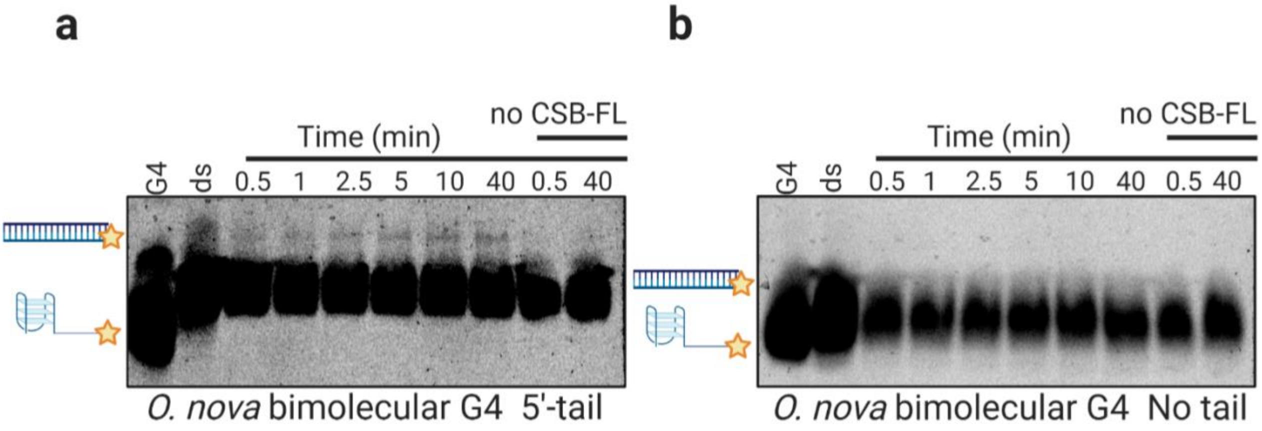


**Figure S10. CSB selectively resolves bimolecular G4s.** Helicase assay gel in presence or absence of 10 nM CSB-FL after 0.5 to 40 min incubation time. (**a**) Helicase assay using 5'-tail *O. nova* G4. (**b**) Helicase assay using untailed *O. nova* G4.

| NAME | SEQUENCE (5'→3') |
|---|---|
| (Cy5) rDNA 5'-tail | ATAATTATAAATAAATAATGGGGCCGGGGGTGGGGTCGGCGGGGAAA |
| (Cy5) rDNA 3'-tail | AAAGGGGCCGGGGGTGGGGTCGGCGGGGATAATTATAAATAAATAATA |
| (Cy5) rDNA-2 5'-tail | ATAATTATAAATAAATAATAGGGTCGGGGGGTGGGGCCCGGGCCGGGG |
| (Cy5) rDNA-3 5'-tail | ATAATTATAAATAAATAATAGGGAGGGAGACGGGGGGG |
| (Cy5) rDNA No tail | GGGGCCGGGGGTGGGGTCGGCGGGGAAA |
| (Cy5) *O. nova* G4 5'-tail | ATAATTATAAATAAATAATGGGGTTTTGGGG |
| (Cy5) *O. nova* G4 No tail | GGGGTTTTGGGG |
| (Cy5) ssDNA | GGCATAGTGCGTGGGCG |
| rDNA-2 No tail | AGGGTCGGGGGGTGGGGCCCGGGCCGGGG |
| rDNA-3 No tail | AGGGAGGGAGACGGGGGGG |
| (FAM) c-KIT1 5'-tail | ATAATTATAAATAAATAATAGGGAGGGCGCTGGGAGGAGGGAAA |
| (FAM) hTELO 5'-tail | ATAATTATAAATAAATAATGGGTTAGGGTTAGGGTTAGGGAAA |
| (FAM) c-MYC 5'-tail | ATAATTATAAATAAATAATTGGGTGGGTAGGGTGGGTAAA |
| (FAM) c-MYC-mut 5'-tail | ATAATTATAAATAAATAATATTGAGTGTGTGTAGTGTGTGTAAA |
| (FAM) c-MYC 3'-tail | AAATGGGTGGGTAGGGTGGGTATAATTATAAATAAATAATA |
| (FAM) c-MYC No tail | TGAGGGTGGGTAGGGTGGGTAA |
| (FAM) c-KIT1 No tail | TGGGAGGGCGCTGGGAGGAGGG |
| (FAM) HRAS 5'-tail | ATAATTATAAATAAATAATATCGGGTTGCGGGCGCAGGGCACGGGCGAAA |
| Complementary rDNA 5'-tail | TTTCCCCGCCGACCCCACCCCCGGCCCCTATTATTTATTTATAATTAT |
| Complementary c-KIT1 5'-tail | TTTCCCTCCTCCCAGCGCCCTCCCTTATTATTTATTTATAATTAT |
| Complementary hTELO 5'-tail | TTTCCCTAACCCTAACCCTAACCCTATTATTTATTTATAATTAT |
| Complementary c-MYC 5'-tail | TTTACCCACCCTACCCACCCATATTATTTATTTATAATTAT |
| Complementary c-MYC-mut 5'-tail | TTTACACACACTACACACACTCAATATTATTTATTTATAATTAT |
| Complementary c-MYC 3'-tail | TATTATTTATTTATAATTATACCCACCCTACCCACCCATTT |
| Complementary c-MYC No tail | TTACCCACCCTACCCACCCTCA |
| Complementary c-KIT1 No tail | CCCTCCTCCCAGCGCCCTCCCA |
| Complementary rDNA no tail | TTTCCCCGCCGACCCCACCCCCGGCCCC |
| Complementary rDNA 3'-tail | TATTATTTATTTATAATTATCCCCGCCGACCCCACCCCCGGCCCCTTT |
| Complementary HRAS 5'-tail | TTTCGCCCGTGCCCTGCCCTGCGCCCGCAACCCGATATTATTTATTTATAATTAT |
| Complementary *O. nova* G4 5'-tail | CCCCAAAACCCCTATTATTTATTTATAATTAT |
| Complementary *O. nova* G4 No tail | CCCCAAAACCCC |

**Table S1. Sequences of the oligonucleotides used.**

| G4 sequence | CSB-FL (0.5-40 min) | NO CSB-FL (0.5-40 min) |
|---|---|---|
| rDNA 5'-tail | 24.6**** | 8**** |
| rDNA-2 5'-tail | 61.1* | 45.7* |
| cKIT1 5'-tail | 45.1 | 40.6 |
| hTELO 5'-tail | 41.2 | 49.4 |
| HRAS 5'-tail | 69.8 | 67.8 |
| cMYC 5'-tail | 36.3 | 28.3 |
| mut-cMYC 5'-tail | 68.9 | 63.3 |
| cMYC 3'-tail | 52.7 | 59.2 |
| rDNA No tail | 24.1 | 24.6 |
| cMYC No tail | 38.1 | 39.2 |
| cKIT1 No tail | 34.8 | 42.9 |

**% double strand (ds)**

**Table S2**. **Quantification of the helicase assay gels testing a panel of different G4-forming sequences represented as percentage of ds formation in presence or absence of CSB-FL.** All the results were based on the average of three independent experiments. Significance was calculated based on a two-tailed Student's t-test. Asterisks indicate statistical difference at 95% confidence levels (CI) between presence or absence of the proteins. *$p < 0.05$, ****$p < 0.0001$.

% double strand (ds)

| G4 sequence | CSB-HD (0.5-40 min) | NO CSB-HD (0.5-40 min) |
|---|---|---|
| rDNA 5'-tail | 37.6* | 23.4* |
| cKIT1 5'-tail | 30.2 | 28.3 |
| cKIT1 No tail | 74.9 | 74.1 |

**Table S3. Quantification of the helicase assay gels testing a panel of different G4-forming sequences represented as percentage of ds formation in presence or absence of CSB-HD.** All the results were based on the average of three independent experiments. Significance was calculated based on a two-tailed Student's t-test. Asterisks indicate statistical difference at 95% confidence levels (CI) between presence or absence of the proteins. *$p < 0.05$, ****$p < 0.0001$.
